# Optogenetic control of the Bone Morphogenetic Protein signalling pathway through engineered blue light-sensitive receptors

**DOI:** 10.1101/2020.04.27.063073

**Authors:** Paul A. Humphreys, Steven Woods, Christopher A. Smith, Stuart A. Cain, Robert Lucas, Susan J. Kimber

**Affiliations:** Division of Cell Matrix & Regenerative Medicine, Faculty of Biology, Medicine and Health, The University of Manchester, Manchester, UK; Division of Neuroscience & Experimental Psychology, Faculty of Biology, Medicine and Health, The University of Manchester, Manchester, UK

**Keywords:** BMP signalling, cell signalling, chondrocytes, light-oxygen-voltage (LOV)-sensing domain, optogenetics

## Abstract

Bone Morphogenetic Proteins (BMPs) are members of the Transforming Growth Factor β (TGFβ) superfamily and have crucial roles during development; including mesodermal patterning and specification of renal, hepatic and skeletal tissues. *In vitro* developmental models currently rely upon costly and unreliable recombinant BMP proteins that do not enable dynamic or precise perturbation of the BMP signalling pathway. Here, we develop a novel optogenetic BMP signalling system (optoBMP) that enables rapid induction of the canonical BMP signalling pathway through illumination with blue light. We demonstrate the utility of the optoBMP system in multiple human cell lines to initiate signal transduction through phosphorylation and nuclear translocation of SMAD1/5, leading to upregulation of BMP target genes including *Inhibitors of DNA binding ID2* and *ID4*. Furthermore, we demonstrate how the optoBMP system can be used to fine-tune activation of the BMP signalling pathway through variable light stimulation. Optogenetic control of BMP signalling will enable dynamic and high-throughput intervention across a variety of applications in cellular and developmental systems.

## Introduction

The Transforming Growth Factor β (TGFβ) superfamily of signalling molecules is a group of structurally related proteins that perform key cellular regulatory functions throughout development and during tissue homeostasis (Massagué, 2012). TGFβ molecules are dimeric when active and initiate signal transduction through binding to a Type I or II receptor. Ligand binding leads to the formation of a heterotetrameric signalling complex consisting of two Type I and two Type II receptors that assemble depending upon ligand specificity (Chen *et al.*, 1998). In canonical signalling, TGFβ superfamily receptors recruit and activate receptor (R)-SMAD proteins, defined by the presence of a short SSXS activation motif which enables their interaction with Type I receptors (Massagué, 1998).

Despite close structural homology, TGFβ superfamily receptors are defined as TGFβ-like or Bone Morphogenetic Protein (BMP)-like depending upon their interaction with one of two R-SMAD groups; SMAD2/3 in TGFβ-like and SMAD1/5/8 in BMP-like signalling. Phosphorylated R-SMAD proteins form a complex with SMAD4 facilitating their translocation to and accumulation within the nucleus. R-SMAD-SMAD4 complexes bind directly to DNA where they, in conjunction with other regulatory transcription factors, control the expression of an extensive variety of genes despite the relatively small number of upstream pathway components (Massagué, 1998). The ability to elicit a diverse range of cellular responses allows TGFβ and BMP signals to have multiple roles in directing cell fate during development, including the specification of germ layers and subsequent specialisation of a wide variety of cell types. Indeed, many established directed differentiation protocols of human pluripotent stem cells (hPSCs) require the activation or inhibition of TGFβ/BMP signals throughout multiple stages to drive differentiation (Hay *et al.*, 2008; Kelleher *et al.*, 2019; Loh *et al.*, 2016; Oldershaw *et al.*, 2010; T. Wang *et al.*, 2019).

*In vitro* investigative and developmental models primarily rely upon activation or inhibition of signalling pathways to elicit a downstream cellular response; predominantly through the use of soluble recombinant growth factors or synthetic chemical compounds that mimic their activity. However, the use of stimulatory molecules to investigate complex biological systems is limited by cellular receptor expression, molecule instability and lack of precise control over signalling kinetics (Mitchell *et al.*, 2016). Biological tools that enable dynamic perturbation of cellular stimuli can more precisely examine the interplay of signalling factors that occur within complex molecular networks. For example, technological advances in microfluidic- and bioactive biomaterial-based approaches have the capacity to provide increased control of signal inputs but are still limited by their complexity and low throughput (DeFail *et al.*, 2006; Tabata & Lutolf, 2017).

Optogenetic technologies, which enable control of cell signalling and physiology through light, are rapidly expanding in scope beyond initial restrictions to light sensitive channel rhodopsins. Advances in optogenetics have been enabled by the engineering of synthetic photoreceptors that consist of a cell signalling protein associated with a photoreceptor sensory domain (Tichy *et al.*, 2019; Ziegler & Möglich, 2015). Commonly employed photoreceptors include flavoprotein blue light response sensors and red light sensitive phytochromes (Conrad *et al.*, 2014; Levskaya *et al.*, 2009). The light-oxygen-voltage (LOV) flavoprotein family of photoreceptors are particularly attractive in photoreceptor engineering due to their small size and versatility of optogenetic manipulations (Pudasaini *et al.*, 2015). Optogenetic tools that utilise the LOV domain have been used to enable optical control of cellular signalling pathways, gene expression regulation and protein localisation (Grusch *et al.*, 2014; Guntas *et al.*, 2015; X. Wang *et al.*, 2012). Optogenetic approaches enable truly dynamic cellular perturbation through spatio-temporally precise optical stimulation techniques that can be finely tuned through modulation of light wavelength, intensity and frequency.

Precise control over TGFβ signalling has been obtained through various means; including chemical-mediated dimerisation of chimeric receptors, sequestering and release of TGFβ ligands by biomaterials and magnetic induction of TGFβ signalling (Gonçalves *et al.*, 2018; Kim *et al.*, 2015; Luo & Lodish, 1996). In recent years, optogenetics has been utilised by several studies to precisely manipulate TGFβ-like receptors and the downstream canonical pathway (Li *et al.*, 2018; Sako *et al.*, 2016). However, there is still a lack of reports that describe control of related BMP-like signalling, despite the essential roles of BMP family ligands in skeletal, hepatic and renal development. In this study we establish an inducible optogenetic BMP signalling system (optoBMP) that enables rapid manipulation of the downstream BMP pathway through blue light stimulation. Blue light illumination results in activation of SMAD1/5/8 signal transduction which initiates BMP-like transcriptional activity. We demonstrate the dynamic nature of the system through manipulation of transcriptional kinetics with modulation of light irradiance. Finally, we demonstrate the applicability of the optoBMP system in multiple cell lines that will allow future incorporation of the optoBMP system into a variety of cellular applications and investigative models.

## Results and Discussion

### Development of an inducible optogenetic BMP signalling system

Activation of the canonical BMP signalling cascade is controlled by the tetramerisation of Type I and II receptors in the presence of an appropriate ligand. We hypothesised that BMP-like Type I and II receptor heterodimerisation would be sufficient to initiate signalling; as previous work has indicated TGFβ-like Type I and II receptor dimerisation can activate the canonical SMAD signalling pathway (Luo & Lodish, 1996; Sako *et al.*, 2016). Therefore, we constructed two optogenetic BMP-like receptors; optoBMPR1B and optoBMPR2 (***Figure 1A-B***). We fused the aureochrome1 Light Oxygen Voltage (LOV) domain derived from *Vaucheria frigida* (Takahashi *et al.*, 2007) with the intracellular regions of both receptors at the C-terminal end and anchored both to the plasma membrane with a myristoylation motif. The *V. frigida* LOV domain dimerises upon blue light stimulation; thus forcing the Type I and II receptors into close proximity to initiate signal transduction (***Figure 1A***). Such a strategy has previously proved successful in the light-induced dimerisation of TGFβ-like nodal receptors and several receptor tyrosine kinases (Grusch *et al.*, 2014; Sako *et al.*, 2016). We then incorporated both receptors into lentiviral doxycycline-inducible vectors to enable transgenic genomic integration combined with temporal control over receptor expression (***Appendix Figure S1A-B***).

**Figure 1 –.**
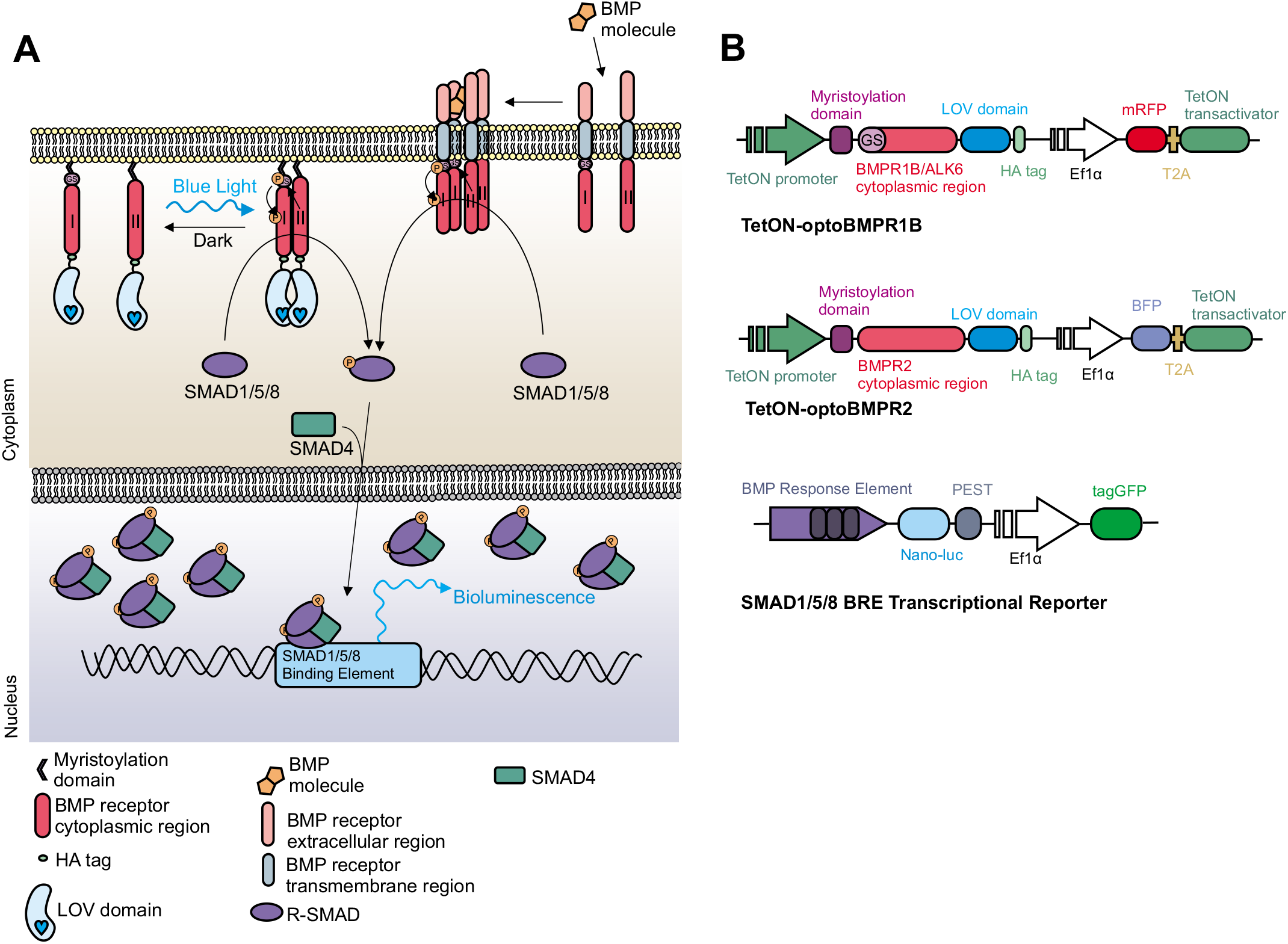
Development of opto-BMP system. (**A**) Schematic representation of the optogenetic BMP (optoBMP) system. (**B**) Vector illustrations of the (**TOP**) optogenetic BMP receptors and (**BOTTOM**) SMAD1/5/8 response element reporter (BRE). Optogenetic receptors were designed as follows: the cytoplasmic region of either BMPR1B/ALK6 or BMPR2 was inserted between a myristoylation signal peptide and the Light Oxygen Voltage (LOV) domain. A haemagglutinin (HA) tag is located at the C-terminus. Optogenetic receptors were then inserted into a doxycycline inducible 2nd generation lentiviral backbone vector with distinct fluorescent protein markers.

### OptoBMP can be used to modulate SMAD1/5/8 transcriptional kinetics

We initially characterised activation of the optoBMP system in HEK293T cells through analysis of real-time SMAD1/5/8 transcriptional kinetics using a BMP-like response element reporter (BRE) (***Figure Appendix S1C***). After 24 hours doxycycline treatment to induce expression of the optoBMP system, optoBMP-HEK293T-BRE cells were illuminated with blue light (470nm, 0.25mW/cm^2^) for 15 minutes. After 4 hours, a significant, over 2-fold induction of the BRE above pre-stimulation was observed in illuminated cells whilst cells that were kept in the dark did not appear to respond (***Figure 2A**).* Optical BRE induction was not significantly different from that obtained with BMP2. Furthermore, we were able to correlate BRE response amplitude with variable light irradiance, demonstrating the ability to fine-tune the optoBMP response with blue light (***Figure 2B**).* Analysis of real-time SMAD1/5/8 transcriptional kinetics demonstrated a unique light-driven response profile in comparison to BMP2-stimulated controls (***Figure 2C***). BMP2 or light illumination resulted in a response peak after ~4 hours, but light illumination resulted in a significantly more rapid induction of the BRE, as illustrated at 30 minutes after initial stimulation (***Figure 2D**).*

**Figure 2 –.**
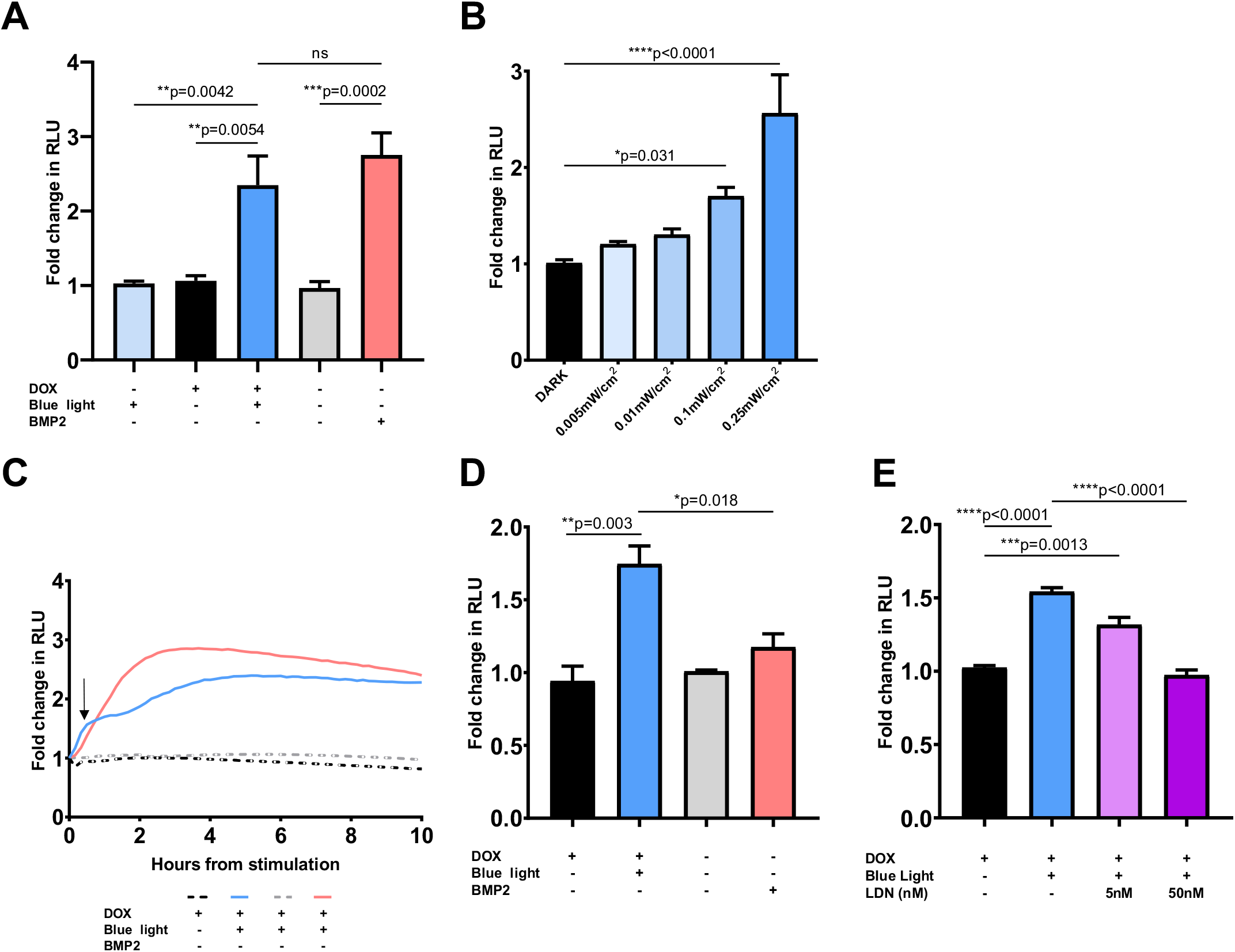
Characterisation of opto-BMP system in optoBMP-HEK293T-BRE cells. **(A)** Analysis of BRE induction 4 hours after stimulation with blue light or 50ng/ml BMP2. N = three wells per condition across five independent experiments. (**B**) Manipulation of BRE induction through stimulation with variable blue light irradiance. N = three wells per condition across four independent experiments. (**C**) Analysis of BRE kinetics over 10 hours after stimulation with blue light or 50ng/ml BMP2. Arrow indicates response after 30 minutes shown in (**D**). N = three wells per condition across three independent experiments. (**D**) Analysis of BRE induction 30 minutes after stimulation with blue light or 50ng/ml BMP2. N = three wells per condition across three independent experiments. (**E**) Analysis of BRE induction 30 minutes after stimulation with blue light and addition of LDN193189. N = two wells per condition across three independent experiments.

**Data information**: In A-E data is presented as mean fold change in NanoLuc Luciferase activity (RLU) from pre-stimulation across independent experiments. In A, B, D and E bars represent mean values + standard error of the mean (SEM). *P* values were generated using an ordinary one-way ANOVA (*p<0.05, **p<0.01, ***p<0.005, ****p<0.0001, ns = not significant).

To ensure the optoBMP system functioned through BMP-like receptor activation, we treated optoBMP-HEK293T-BRE cells with LDN193189; which directly inhibits the kinase activity of BMP-like Type I receptors (Cuny *et al.*, 2008). Addition of LDN193189 at a low concentration (5nM) is sufficient to inhibit BMPR1A/ALK3 kinase activity but not that of BMPR1B/ALK6, which we utilised in the design of the optoBMP system (Sanvitale *et al.*, 2013). We observed significant BRE induction in cells treated with 5nM LDN193189 30 minutes after light illumination in comparison to cells that remained in the dark (***Figure 2E**).* Upon application of 50nM LDN193189, BRE induction was abolished, indicating that the optoBMP system initiates canonical BMP-like signal transduction through light-induced opto-BMPR1B kinase activity.

Through targeting the most upstream pathway components for optical control, the optoBMP system almost entirely utilises native cellular machinery. As the binding of SMAD1/5/8 to transcriptional targets are influenced and directed by cell-dependent expressed cofactors, the design of the optoBMP system does not interfere with this process. Available synthetic compounds that target the BMP pathway subvert receptor activation and thus may disrupt additional recruitment of signalling effectors that bind to active SMAD1/5/8 (Genthe *et al.*, 2017; Bradford *et al.*, 2019). Therefore, the optoBMP system has greater potential to investigate the roles of BMP signals in any natural cellular context. As expression and activation of the optoBMP system can be induced at any time upon doxycycline and light illumination respectively, the system subverts native cellular regulatory systems driving the expression of specific receptors or the internalisation and degradation of ligand-receptor complexes (Huang and Chen, 2012). In addition, BMP ligands are promiscuous in their receptor activation and thus light illumination provides precise dosage and downstream receptor specificity.

### Blue light stimulation of optoBMP drives a BMP-like response in a chondrogenic cell line

To investigate the potential of the optoBMP system in a relevant context, we transferred the system into the immortalised chondrocyte cell line TC28a2 to test the response in a skeletal model (Goldring *et al.*, 1994). BMP signals are critical in chondrocyte and osteocyte development and maintenance (Salazar *et al.*, 2016). Optogenetic receptor expression (detected through the tagged haemagglutinin (HA) signal) was only seen in doxycycline treated cells (***Figure 3A**),* confirming the doxycycline-dependent induction of optogenetic receptor expression. We then investigated if activation of the optoBMP system induced native SMAD1/5/8 signal transduction. Nuclear accumulation of P-SMAD1/5 was analysed 2 hours after 15 minutes blue light illumination (***Figure 3B**).* Single cell quantification of the mean nuclear intensity of P-SMAD1/5 indicated a significant increase in comparison to controls that remained in dark conditions (***Figure EV1A**).* P-SMAD1/5 nuclear intensity was observed to be higher in BMP2-treated controls that were stimulated continuously for 2 hours, but the percentage of positive cells in both light and BMP2-stimulated conditions after 2 hours were not significantly different (***Figure 3C**).*

**Figure 3 –.**
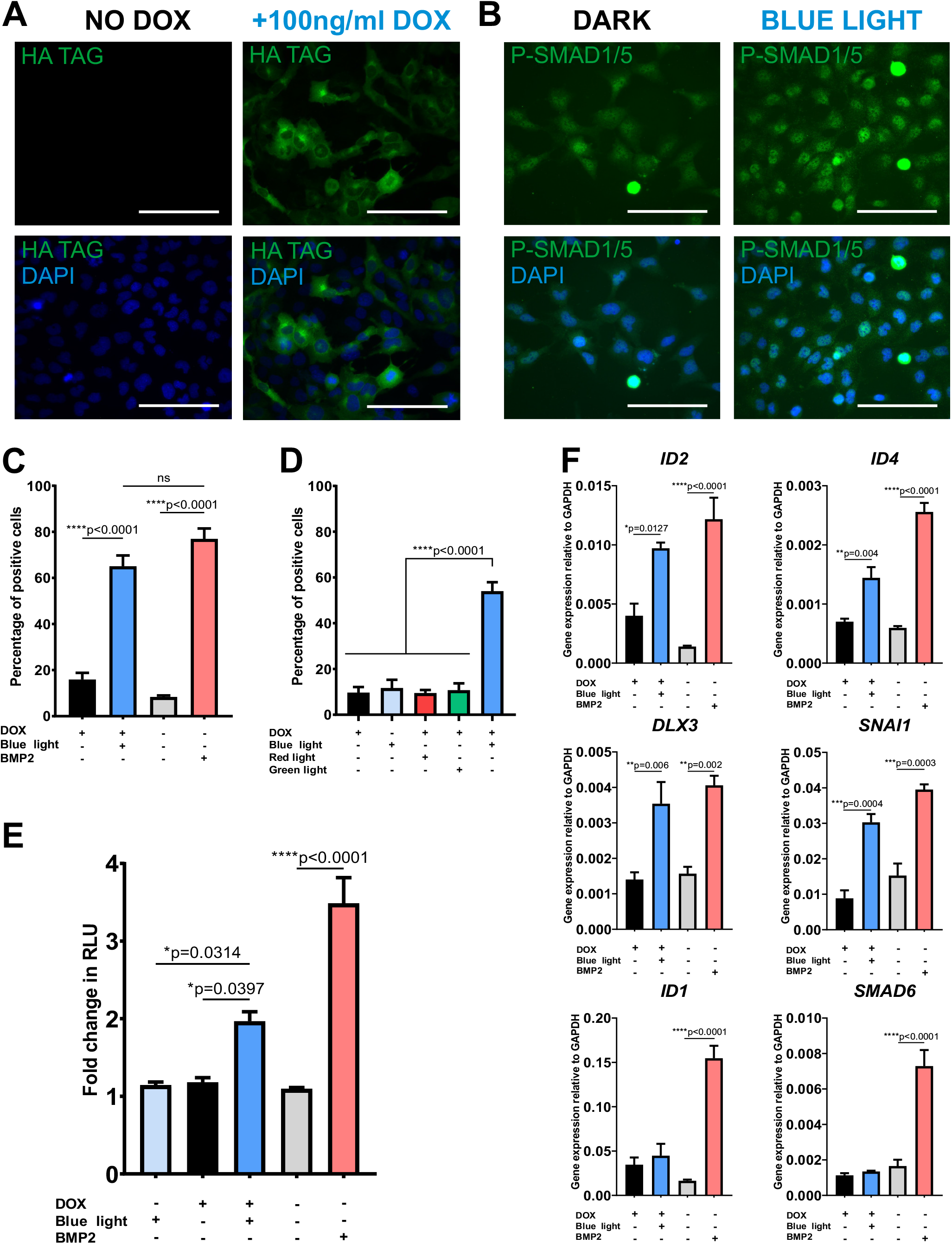
Induction of a BMP-like response in chondrogenic optoBMP-TC28a2 cells. (**A**) Representative immunofluorescence images of cells stained for HA tag with or without addition of 100ng/ml doxycycline for 24 hours. (**B**) Representative immunofluorescence images of cells stained for P-SMAD1/5 2 hours after 15 minutes blue light illumination (**RIGHT**) or having been kept in the dark (**LEFT**). (**C**) Percentage of P-SMAD1/5 positive cells calculated through single-cell quantification of mean nuclear P-SMAD1/5 fluorescence intensity. Cells were stimulated with blue light illumination or 50ng/ml BMP2. Threshold for positivity calculated through analysis of unstimulated controls. N = three different fields of view per condition across three independent experiments. (**D**) Percentage of P-SMAD1/5 positive cells when stimulated with variable light wavelengths. Cells were stimulated with either red, blue or green light (0.25mW/cm^2^) or kept in the dark. N = three different fields of view per condition across three independent experiments. (**E**) Analysis of BRE induction in optoBMP-TC28a2-BRE cells 4 hours after stimulation with blue light or 50ng/ml BMP2. Bars represent mean fold change in RLU value 4 hours from pre-stimulation +SEM. N = three wells per condition across three independent experiments. (**F**) Gene expression analyses of BMP-target genes. Cells were stimulated with blue light illumination or 50ng/ml BMP2 and analysed after 4 hours. Gene expression was normalised to GAPDH. N = four independent experiments. **Data information**: Scale bars in A-B represent 100μm. Data presented in C-F represent mean values + SEM. *P* values were generated using an ordinary one-way ANOVA (*p<0.05, **p<0.01, ***p<0.0005, ****p<0.0001, ns = not significant).

We observed a small increase in basal SMAD1/5/8 signalling upon doxycycline addition in the absence of the light; indicating a low level of spontaneous optoBMP activation, most likely as a result of the spatial proximity of receptors both expressed at the cell membrane. Alternatively, or in combination, continuous doxycycline-driven receptor expression may result in receptor excess. However, blue light illumination resulted in significant enhancement of SMAD1/5 phosphorylation and induction of transcriptional activity in comparison to cells kept in the dark. Although not an issue in this study, spontaneous optoBMP activation without light may be eliminated through a similar approach to that used by Li *et al*. (2018); in which a single receptor is localised to the cytoplasm, hence allowing signalling solely thorough light-activated membrane localisation. The blue light specificity of the optoBMP system was then demonstrated through subjecting optoBMP-TC28a2 cells to variable light wavelength stimulations (***Figure EV1B***). There was no significant difference in mean percentage of cells positive for nuclear P-SMAD1/5 between cells left in the dark and cells illuminated with red (670nm) or green (560nm) light (***Figure 3D**).* In addition, doxycycline addition, and therefore expression of the optoBMP system, was required to enable blue-light induction of P-SMAD1/5 nuclear accumulation.

To illustrate the ability of the optoBMP system to elicit a BMP-like transcriptional response in optoBMP-TC28a2 cells, we initially measured the SMAD1/5/8 response through BRE induction. As with optoBMP-HEK293T cells, we found that 15 minutes of blue light illumination resulted in significant induction of a SMAD1/5/8 response after 4 hours (***Figure 3E**).* We then investigated whether BRE induction correlated with upregulation of direct BMP-pathway target genes 4 hours after blue light illumination. Blue light induction of optoBMP resulted in significant upregulation of *Inhibitor of Differentiation 2 (ID2), ID4, Distal-Less Homeobox 3 (DLX3*) and *Snail Family Transcriptional Repressor 1 (SNAI1*) which are all known to be downstream targets of SMAD1/5/8 (***Figure 3F***) (Hollnagel *et al.*, 1999; Savary *et al.*, 2013; Yang *et al.*, 2014). However, blue light illumination did not lead to upregulation of known BMP response genes *ID1* or inhibitory *SMAD6,* which acts as part of a negative feedback loop, both of which are upregulated following BMP2 stimulation (Hata *et al.*, 1998). The apparent differences between optoBMP- and BMP2 ligand-driven responses may be as a result of unique signalling receptor complexes. The lack of native BMPR1B/ALK6 expression in TC28a2 cells indicates a BMP signalling response is driven through a BMPR1A/ALK3-BMPR2 receptor complex unlike the optoBMP system (***Figure EV2**).* Alternatively, BMP2 stimulation used here involved 4 hours of sustained signalling in comparison to a short 15 minute light-induced activation of the BMP pathway, which may be mimicked in the future through using longer durations of light illumination.

### Application of optoBMP in a human development model

As human embryonic stem cells (hESCs) have proved a useful model system for studying development to many different lineages, they are a suitable system for testing optogenetic regulation of development. To demonstrate the potential of the optoBMP system in this context we therefore transduced hESC line MAN13 (Ye *et al.*, 2017) with the optoBMP system. We initially performed verification of pluripotency through immunofluorescent staining of established pluripotent markers OCT4 and NANOG (***Figure 4A**).* Then, to determine functionality of the optoBMP system in hESCs, we analysed nuclear accumulation of P-SMAD1/5 after blue light illumination (***Figure 4B**).* Single cell quantification of nuclear P-SMAD1/5 indicated significant induction of the canonical BMP pathway when compared to cells kept in the dark (***Figure 4C**).* To further illustrate optogenetic activation in hESCs we analysed BMP target gene expression. Again we observed upregulation of direct BMP target genes including *ID2*, *ID3*, *ID4 and DLX3 (**Figure EV3**).* Interestingly, several genes appeared upregulated by light illumination and not by BMP2, including differentiation genes *SNAI1* and *SOX9.* As *ID, SNAIL* and *SOX* genes have crucial roles during developmental processes, these preliminary data suggest that the optoBMP system could be utilised in hPSC development models as a substitute for costly recombinant growth factor supplementation (Evseenko *et al.*, 2010; Hollnagel *et al.*, 1999). Modification of the optoBMP expression system through CRISPR/Cas9-mediated incorporation into genomic safe harbour locus AAVS1 would be useful in the future to enable uniform and ubiquitous expression without lentiviral transgene silencing; an issue which has been commonly reported in the differentiation of pluripotent stem cells (Benabdellah *et al.*, 2014; Herbst *et al.*, 2012; Pfaff *et al.*, 2013).

**Figure 4 –.**
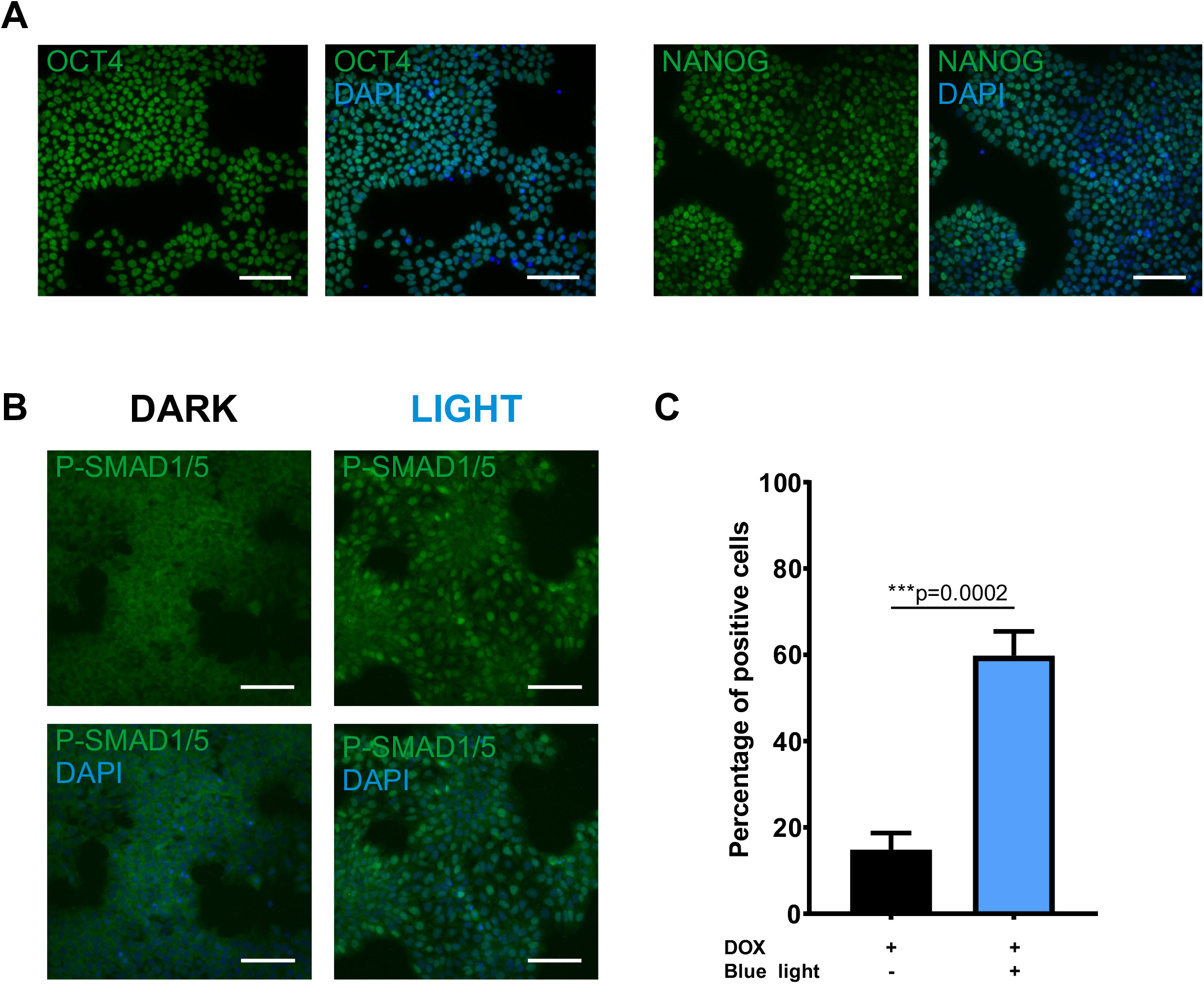
Optogenetic activation of canonical BMP signalling in pluripotent optoBMP-hESCs. **(A)** Verification of pluripotency. Representative immunofluorescence images of optoBMP-hESCs stained for pluripotent markers OCT4 and NANOG. **(B)** Representative immunofluorescence images of cells stained for P-SMAD1/5 2 hours post 15 minutes blue light illumination. **(C)** Percentage of P-SMAD1/5 positive cells calculated through single-cell quantification of mean nuclear P-SMAD1/5 intensity. Threshold for positivity calculated through analysis of unstimulated controls. N = three different fields of view per condition across three independent experiments **(D)** Gene expression analyses of BMP-target genes. Cells were stimulated with 15 minutes blue light illumination or 50ng/ml BMP2. Gene expression was normalised to GAPDH. N = two independent experiments. **Data information**: Scale bars in A-B represent 100μm. Data presented in C represent mean values + SEM. *P* value was generated using an un-paired two-tailed students *t*-test (***p<0.0005).

In this study we demonstrate that short light pulses are sufficient to alter BMP signalling using the optoBMP system, although in the future it will be interesting to investigate the effect of complex light stimulation patterns and longer illumination periods as used in some other optogenetic studies (Gerhardt *et al.*, 2016; Li *et al.*, 2018; Repina *et al.*, 2019). Furthermore, sustained light illumination is most likely required for driving complex cellular responses such as in directing cell fate decisions and differentiation, as illustrated by the recent application of an optogenetic Wnt tool in hPSCs (Repina *et al.*, 2019). In summary, we report the development of a novel optogenetic signalling system that enables control of canonical BMP-like SMAD1/5/8 signal transduction through blue light-sensitive BMP receptors. We demonstrate that the optoBMP system can reliably recapitulate initiation of the canonical BMP signalling cascade in several cell lines through phosphorylation and nuclear translocation of SMAD1/5/8 leading to BMP-like transcriptional activation, including upregulation of direct BMP gene targets. Extension of the optogenetic toolkit to target the BMP pathway will provide the means to dynamically probe BMP signals in a wide variety of systems.

## Materials & Methods

### DNA vector assembly

All vectors were constructed through NEB HiFi Assembly or restriction enzyme cloning. Both optogenetic receptors were assembled through an initial generation of a vector backbone through PCR amplification of opto-mFGFR (a gift from Harald Janovjak, Addgene plasmid #58745) to omit the mFGFR coding region. The intracellular coding regions of human BMPR1B/ALK6 (fully sequenced cDNA clone from Source Bioscience; vector #IRATp970H1175D) and BMPR2 (Source Bioscience, #IRATp970B1178D) identified from UniProt (https://www.uniprot.org/) were then PCR amplified and inserted into the vector backbone through NEB HiFi Assembly (New England Biolabs, #E2621). Each optogenetic receptor was then subsequently inserted into a second-generation lentiviral shuttle doxycycline inducible vector through NEB HiFi Assembly. Briefly, vector inserts were generated through PCR amplification of each optogenetic receptor with 30bp overhangs. Vector backbones, pCDH-TRE3G-MCS-EF1a-tagRFP-T2A-TetON3G or pCDH-TRE3G-MCS-EF1a-tagBFP-T2A-TetON3G for Type I or II receptor respectively, were then digested with EcoRI and vector inserts were annealed in through NEB HiFi Assembly. Final lentiviral shuttle vectors were designated TetOn-optoBMPR1B and TetOn-optoBMPR2.

A modified version of a BMP reporter vector previously described (Lockhart-Cairns *et al.*, 2019) was constructed. The BMP response element (BRE) consisted of multiple SMAD binding elements arranged in tandem in forward and reverse orientations, placed upstream of the AAV minimal late promoter (Korchynskyi & Ten Dijke, 2002). The BRE was cloned upstream of a destabilized form of NanoLuc Luciferase (Promega) containing a C-terminal protein degradation sequence (PEST sequence, NLucP). The completed BRE-NLucP was cloned into a modified version of the lentiviral expression vector pCDH-EF1-T2A-copGFP, resulting in the construct pCDH-BRE-nLUCP-EF1a-copGFP. Vector maps were generated using Snapgene software (***Figure Appendix S1***).

### Cell culture

HEK293T (ATCC, #CRL-11268), TC28a2 (Goldring *et al.*, 1994) and SW1353 (ATCC, #HTB-94) cell lines were cultured in 75cm^2^ cell culture flasks (Corning) with DMEM (Gibco, #11960044) containing 10% w/v fetal bovine serum (Merck, #12103C), 1% w/v L-glutamine (Gibco, #25030081) and 1% w/v penicillin/streptomycin (Gibco, #15140122). For routine maintenance cells were sub-cultured into flasks containing fresh warmed medium at a passage ratio of 1:10. Briefly, cells were washed with Phosphate Buffered Saline (PBS – Merck, #D8537) before cell dissociation with 3ml TryPLE Express solution (Gibco, #12604021). Cells were then centrifuged at 700xg for 3 minutes before pellet resuspension and continued passage.

MAN13 human embryonic stem cells (hESCs) (Ye *et al.*, 2017) were cultured in 6-well plates (Corning) coated with 5μg/ml human recombinant vitronectin (Life Technologies, #A14700) in mTesR1 (StemCell Technologies, #5850) with medium replaced every two days. For routine maintenance culture, cells were washed with PBS before dissociation using 0.5mM EDTA solution (Invitrogen, #15575-038). Cells were centrifuged at 700xg for 3 minutes before subculture into fresh medium containing 1X RevitaCell Supplement (Life Technologies, #A2644501).

### Lentiviral particle production and cell line generation

All lentiviral particles were generated through transfection of HEK293T cells using calcium chloride precipitation. Briefly, HEK293T cells were seeded in 15cm^2^ dishes at 8×10^6^ cells/dish 24 hours before transfection with 9μg psPAX2, 6μg pMD2.G (VSV), 12μg shuttle plasmid (TetOn-optoBMPR1B, TetOn-optoBMPR2 or pCDH-BRE-nLUCP-EF1a-copGFP) per dish. Medium containing lentiviral particles was collected over 48 hours and concentrated at 6000xg overnight at 4°C. The lentiviral pellet was then resuspended in 10ml ice-cold PBS and was centrifuged in a high-speed SW40-Ti rotor (Beckman Coulter) at 50,000xg for 90 minutes at 4°C. Finally, the concentrated pellet was resuspended in 200μl ice-cold PBS and stored at −80°C. Viral titres were calculated through serial dilution of viral preparations before transduction of HEK293T cells and flow cytometry analysis of appropriate fluorescent marker. HEK293T and TC28a2 cells were transduced with viral particles at a multiplicity of infection (MOI) that did not exceed 20 IU/cell, whilst MAN13 cells were transduced at an MOI that did not exceed 10 IU/cell before being sorted through FACS (BD FACS Aria Fusion).

### Optogenetic and chemical stimulation

Prior to optical stimulation experiments, all cell lines were seeded in black-walled 96 well plates at a density of 1×10^4^ cells/well or and were maintained in the dark after induction of optogenetic receptor expression through 24 hours incubation in serum-free medium containing 100ng/ml doxycycline hyclate (Merck, #D9891). Optical stimulation was performed at room temperature using either a custom-built Arduino-controlled LED array device using 470nm LEDs (Ballister *et al.*, 2018) or CoolLED pE4000 at 460nm. Light irradiance was controlled through a piece of custom software (LED array) or directly from the equipment control panel (CoolLED). Light illumination was performed at 0.25mW/cm^2^ for 15 minutes unless otherwise highlighted. Irradiance was measured using a spectroradiometer (SpectroCAL MKII; Cambridge Research Systems). Alongside optogenetic stimulation where noted, cells were stimulated with 50ng/ml BMP2 (R&D Systems, #355-GMP). Inhibition of the BMP signalling pathway was achieved through addition of LDN193187 dihydrochloride (Tocris, #6053) in combination with optical stimulation at concentrations noted. Alongside LDN193187, cells were treated with DMSO vehicle control.

### Nano-Glo© luciferase assay system

Optogenetic HEK293T and TC28a2 cells were seeded in black-walled 96-well plates before 24 hours serum starvation and induction of optogenetic receptor expression. Extended NanoLuc luciferase substrate Vivazine™ (Promega, #N2580) was added to live cells (dilution to 1X in serum-free DMEM) 2 hours prior to stimulations. After performing optogenetic or chemical stimulations, expression of NanoLuc luciferase was detected in an Alligator Luminescence System (Cairn Research) using an appropriate exposure time (10-30 minutes). Bioluminescent micrographs taken over a period of time were stacked using ImageJ software. A region of interest (ROI) was then drawn around each well and relative luminescent units (RLU) were calculated for each micrograph through Z-stack analysis. Luminescence was normalised to background and to luminescence value prior to the start of stimulation.

### Immunocytochemistry and imaging

Cells were fixed in 4% paraformaldehyde in PBS for 15 minutes at room temperature and subsequently washed in PBS three times. Fixed cells were then permeabilised and blocked in 0.1% Triton-X (Merck, #9002-93-1), 10% donkey serum (DS) (Merck, #D9663) in PBS for 30 minutes at room temperature, before a further three PBS washes. Cells were incubated with primary antibodies (Cell Signalling Technologies; #3724 HA tag 1:700 dilution, #9516 P-SMAD1/5 1:200 dilution, #2890 OCT4-A 1:200 dilution, #4903 NANOG 1:200 dilution) in 1% DS in PBS at 4°C overnight before being subsequently washed three times in PBS. Cells were then treated with AlexaFluor™-488- or Alexa-Fluor™-594-labelled secondary antibodies (Life Technologies #A32790/#A32754, 1.200 dilution in PBS 1% DS) for 45 minutes at room temperature before a final three PBS washes. Nuclei were visualised through staining with 10μg/ml 4,6-diamindino-2-phenylindole (DAPI - Life Technologies, #D1306) in PBS for 5 minutes at room temperature followed by three washes in PBS. Fluorescent micrographs were taken using a Zeiss Axioimager D2 upright microscope and captured using a Coolsnap HQ2 camera (Photometrics) through micromanager software v1.4.23.

### Image analysis

Image processing and quantification was performed using ImageJ software. Nuclear quantification of P-SMAD1/5 was performed through an initial identification of nuclear outlines through the DAPI channel (358nm). This was achieved through the ‘Analyse Particles’ function of ImageJ after converting the image to a binary output. Each nuclear outline identified by ImageJ was then applied to the P-SMAD1/5 channel (488nm) before using the ROI manager to measure the mean grey value within each nuclear outline. Values that fell 2*SD from the mean were omitted from analysis to eliminate false positives or negatives. A threshold mean intensity value to determine a positive P-SMAD1/5 response was calculated from analysis of non-stimulated control images.

### RNA extraction, reverse transcription and qPCR analysis

RNA extraction and purification was performed using a Qiagen RNeasy mini-kit (Qiagen, #74104) according to the manufacturer’s instructions. Briefly, cells were dissociated and the cell pellet was resuspended in 350μl RLT lysis buffer. Cell lysate was then subjected to a series of on-column washes, after addition of 350μl 70% ethanol, before elution in nuclease-free water (Invitrogen, #4387937). After quantification of RNA concentration, 2μg RNA was converted to cDNA using a high capacity cDNA reverse transcription kit (Thermo Fisher Scientific, #4368813). qPCR reaction was then prepared using PowerUp SYBR green master mix (ThermoFisher Scientific, #A25742) with 10ng cDNA per reaction and 400nM final concentration forward and reverse primers. qPCR reaction was ran using a BioRad C1000Touch™ Thermal Cycler using the following cycling conditions: denaturation at 95°C for 10 minutes, 39 cycles of 95°C for 30 seconds, 60°C for 30 seconds and 72°C for 35 seconds, final extension at 72°C for 10 minutes and melt curve analysis at 65°C for 5 seconds and 95°C for 30 seconds. Raw qPCR data was normalised to GAPDH using the 2^-ΔCT^ method. Primer sequences are listed in ***Table 1***.

**Table 1 –.**
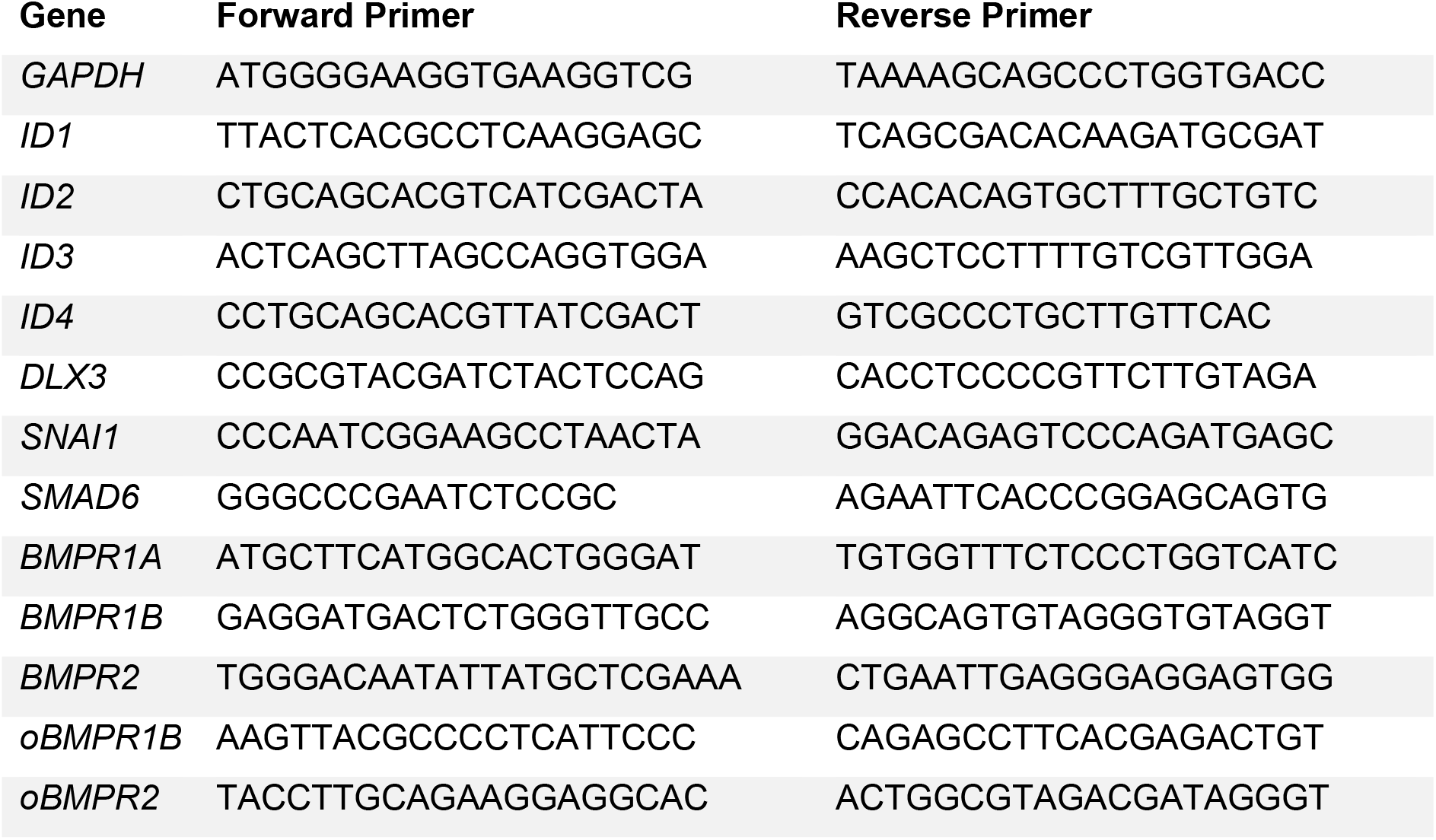
List of primer sequences used for qPCR reactions. oBMPR1B and oBMPR2 indicate synthetic optogenetic transgenes.

### Statistical Analyses and Figure Generation

Figures were generated using CorelDraw Graphics Suite or GraphPad Prism 8 software. Statistical analyses were performed using GraphPad Prism 8. Comparison between two groups were performed using an unpaired two-tailed student’s *t*-test and more than two groups with an ordinary one-way analysis of variance (ANOVA) and subsequent Tukey’s multiple comparisons test. A *p* value of <0.05 was indicative of significance.

## Acknowledgements

The authors acknowledge financial support from the Engineering and Physical Sciences Research Council (EPSRC) and the Medical Research Council (MRC): Centre for Doctoral Training (CDT) in Regenerative Medicine (EP/L014904/1) studentship to PH with additional support from Arthritis Research UK (Grant R20786) and Medical Research Council (Grant MR/S002553/1). TC28a2 cells were a gift from Dr L. Reynard (Newcastle University). We are grateful to Nicola Bates for technical support.

## Author Contributions

Project conception by SJK and RL. Experimental design by SJK and PH. PH performed all experiments and data analysis. SC generated lentiviral TetON plasmids and BRE reporter plasmid. SW generated optoBMP-hESC line. Data interpretation by PH, SW, CAS, RL and SJK. Manuscript written by PH, SW, CAS and SJK.

## Expanded and Supplementary Figure Legends

**Appendix Figure S1 - Vector maps.**

**(A)** Full vector map of TetOn-optoBMPR1 B

**(B)** Full vector map of TetOn-optoBMPR2

**(C)** Full vector map of pCDH-BRE-nLUCP-EF1a-copGFP

**Figure EV1 - Single cell quantification of nuclear P-SMAD1/5 fluorescence.**

**(A-B)** Quantification of nuclear P-SMAD1/5 mean grey value when (**A**) cells were stimulated with blue light or 50ng/ml BMP2 and (**B**) stimulated with a range of light wavelengths. N = three different fields of view per condition across three independent experiments.

**Data Information**: Nuclei were identified with DAPI channel and relative nuclear fluoresence intensity of P-SMAD1/5 was measured with ImageJ. Quantification performed using three different fields of view per condition across three independent experiments. Violin plots represent all quantified data. Dotted red line indicates mean grey value threshold for positivity determined through analysis of non-stimulated controls. *P* values were generated using an ordinary one-way ANOVA (****p<0.0001).

**Figure EV2 - Gene expression analyses of BMP-like receptors in optoBMP-TC28a2 cells.**

**Data Information:** Native *BMPR1B* and *BMPR2* transcript was amplified through primer design targeting exons within the extra-cellular coding regions. *BMPR1B* transcript was not detected (nd) and primers were validated in an alternative chondrosarcoma SW1353 cell line. Optogenetic receptor transcript was detected through reverse direction primer design targeting the LOV domain coding region. Gene expression was normalised to GAPDH. N = four independent experiments. Bars represent mean values + SEM. *P* values were generated using an ordinary one-way ANOVA (*p<0.05, **p<0.01, ****p<0.0001).

**Figure EV3 - Gene expression analyses of BMP-target genes in pluripotent optoBMP-hESCs**. Cells were stimulated with 15 minutes blue light illumination or 50ng/ml BMP2 and analysed after 4 hours. Gene expression was normalised to GAPDH. N = two independent experiments.

## References

Ballister, E. R., Rodgers, J., Martial, F., & Lucas, R. (2018). A live cell assay of GPCR coupling allows identification of optogenetic tools for controlling Go and Gi signaling. BMC Biology, 16(1), 1–16. https://doi.org/10.1186/s12915-017-0475-2

Benabdellah, K., Gutierrez-Guerrero, A., Cobo, M., Muñoz, P., & Martín, F. (2014). A chimeric HS4-SAR insulator (IS2) that prevents silencing and enhances expression of lentiviral vectors in pluripotent stem cells. PLoS ONE, 9(1). https://doi.org/10.1371/journal.pone.0084268

Chen, Y.-G., Hata, A., Lo, R., Wotton, D., Shi, Y., Pavletich, N., & Massague, J. (1998). Determinants of specificity in TGF-β signal transduction. Genes & Development, 12, 2144–2152. https://doi.org/10.1101/gad.12.14.2144

Conrad, K. S., Manahan, C. C., & Crane, B. R. (2014). Photochemistry of flavoprotein light sensors. Nature Chemical Biology, 10(10), 801–809. https://doi.org/10.1038/nchembio.1633

Cuny, G. D., Yu, P. B., Laha, J. K., Xing, X., Liu, J. F., Lai, C. S., Deng, D. Y., Sachidanandan, C., Bloch, K. D., & Peterson, R. T. (2008). Structure-activity relationship study of bone morphogenetic protein (BMP) signaling inhibitors. Bioorganic and Medicinal Chemistry Letters, 18(15), 4388–4392. https://doi.org/10.1016/j.bmcl.2008.06.052

DeFail, A. J., Chu, C. R., Izzo, N., & Marra, K. G. (2006). Controlled release of bioactive TGF-β1 from microspheres embedded within biodegradable hydrogels. Biomaterials, 27(8), 1579–1585. https://doi.org/10.1016/j.biomaterials.2005.08.013

Evseenko, D., Zhu, Y., Schenke-Layland, K., Kuo, J., Latour, B., Ge, S., Scholes, J., Dravid, G., Li, X., MacLellan, W. R., & Crooks, G. M. (2010). Mapping the first stages of mesoderm commitment during differentiation of human embryonic stem cells. Proceedings of the National Academy of Sciences of the United States of America, 107(31), 13742–13747. https://doi.org/10.1073/pnas.1002077107

Gerhardt, K. P., Olson, E. J., Castillo-Hair, S. M., Hartsough, L. A., Landry, B. P., Ekness, F., Yokoo, R., Gomez, E. J., Ramakrishnan, P., Suh, J., Savage, D. F., & Tabor, J. J. (2016). An open-hardware platform for optogenetics and photobiology. Scientific Reports, 6(June), 1–13. https://doi.org/10.1038/srep35363

Goldring, M. B., Birkhead, J. R., Suen, L. F., Yamin, R., Mizuno, S., Glowacki, J., Arbiser, J. L., & Apperley, J. F. (1994). Interleukin-1 beta-modulated gene expression in immortalized human chondrocytes. J Clin Invest, 94(6), 2307–2316.

Gonçalves, A. I., Rotherham, M., Markides, H., Rodrigues, M. T., Reis, R. L., Gomes, M. E., & El Haj, A. J. (2018). Triggering the activation of Activin A type II receptor in human adipose stem cells towards tenogenic commitment using mechanomagnetic stimulation. Nanomedicine: Nanotechnology, Biology, and Medicine, 14(4), 1149–1159. https://doi.org/10.1016/j.nano.2018.02.008

Grusch, M., Schelch, K., Riedler, R., Reichhart, E., Differ, C., Berger, W., Inglés-Prieto, Á., & Janovjak, H. (2014). Spatio-temporally precise activation of engineered receptor tyrosine kinases by light. The EMBO Journal, 33(15), 1713–1726. https://doi.org/10.15252/embj.201387695

Guntas, G., Hallett, R. A., Zimmerman, S. P., Williams, T., Yumerefendi, H., Bear, J. E., & Kuhlman, B. (2015). Engineering an improved light-induced dimer (iLID) for controlling the localization and activity of signaling proteins. Proceedings of the National Academy of Sciences of the United States of America, 112(1), 112–117. https://doi.org/10.1073/pnas.1417910112

Hata, A., Lagna, G., Massagué, J., & Hemmati-Brivanlou, A. (1998). Smad6 inhibits BMP/Smad1 signaling by specifically competing with the Smad4 tumor suppressor. Genes and Development, 12(2), 186–197. https://doi.org/10.1101/gad.12.2.186

Hay, D. C., Fletcher, J., Payne, C., Terrace, J. D., Gallagher, R. C. J., Snoeys, J., Black, J. R., Wojtacha, D., Samuel, K., Hannoun, Z., Pryde, A., Filippi, C., Currie, I. S., Forbes, S. J., Ross, J. A., Newsome, P. N., & Iredale, J. P. (2008). Highly efficient differentiation of hESCs to functional hepatic endoderm requires ActivinA and Wnt3a signaling. Proceedings of the National Academy of Sciences of the United States of America, 105(34), 12301–12306. https://doi.org/10.1073/pnas.0806522105

Herbst, F., Ball, C. R., Tuorto, F., Nowrouzi, A., Wang, W., Zavidij, O., Dieter, S. M., Fessler, S., Van Der Hoeven, F., Kloz, U., Lyko, F., Schmidt, M., Von Kalle, C., & Glimm, H. (2012). Extensive methylation of promoter sequences silences lentiviral transgene expression during stem cell differentiation in vivo. Molecular Therapy, 20(5), 1014–1021. https://doi.org/10.1038/mt.2012.46

Hollnagel, A., Oehlmann, V., Heymer, J., Rüther, U., & Nordheim, A. (1999). Id genes are direct targets of bone morphogenetic protein induction in embryonic stem cells. Journal of Biological Chemistry, 274(28), 19838–19845. https://doi.org/10.1074/jbc.274.28.19838

Kelleher, J., Dickinson, A., Cain, S., Hu, Y., Bates, N., Harvey, A., Ren, J., Zhang, W., Moreton, F. C., Muir, K. W., Ward, C., Touyz, R. M., Sharma, P., Xu, Q., Kimber, S. J., & Wang, T. (2019). Patient-Specific iPSC Model of a Genetic Vascular Dementia Syndrome Reveals Failure of Mural Cells to Stabilize Capillary Structures. Stem Cell Reports, 13(5), 817–831. https://doi.org/10.1016/j.stemcr.2019.10.004

Kim, J., Lin, B., Kim, S., Choi, B., Evseenko, D., & Lee, M. (2015). TGF-β1 conjugated chitosan collagen hydrogels induce chondrogenic differentiation of human synovium-derived stem cells. Journal of Biological Engineering, 9(1), 1–11. https://doi.org/10.1186/1754-1611-9-1

Korchynskyi, O., & Ten Dijke, P. (2002). Identification and functional characterization of distinct critically important bone morphogenetic protein-specific response elements in the Id1 promoter. Journal of Biological Chemistry, 277(7), 4883–4891. https://doi.org/10.1074/jbc.M111023200

Levskaya, A., Weiner, O. D., Lim, W. A., & Voigt, C. A. (2009). Spatiotemporal control of cell signalling using a light-switchable protein interaction. Nature, 461(7266), 997–1001. https://doi.org/10.1038/nature08446

Li, Y., Lee, M., Kim, N., Wu, G., Deng, D., Kim, J. M., Liu, X., Heo W. Do, & Zi, Z. (2018). Spatiotemporal Control of TGF-β Signaling with Light. ACS Synthetic Biology, 7(2), 443–451. https://doi.org/10.1021/acssynbio.7b00225

Lockhart-Cairns, M. P., Lim, K. T. W., Zuk, A., Godwin, A. R. F., Cain, S. A., Sengle, G., & Baldock, C. (2019). Internal cleavage and synergy with twisted gastrulation enhance BMP inhibition by BMPER. Matrix Biology, 77, 73–86. https://doi.org/10.1016/j.matbio.2018.08.006

Loh, K. M. M., Chen, A., Koh, P. W. W., Deng, T. Z. Z., Sinha, R., Tsai, J. M. M., Barkal, A. A. A., Shen, K. Y. Y., Jain, R., Morganti, R. M. M., Shyh-Chang, N., Fernhoff, N. B. B., George, B. M. M., Wernig, G., Salomon, R. E. E. A., Chen, Z., Vogel, H., Epstein, J. A. A., Kundaje, A., … Weissman, I. L. L. (2016). Mapping the Pairwise Choices Leading from Pluripotency to Human Bone, Heart, and Other Mesoderm Cell Types. Cell, 166(2), 451–467. https://doi.org/10.1016/j.cell.2016.06.011

Luo, K., & Lodish, H. (1996). Signalling by chimeric erythropoietin-TGFB receptors: homodimerization of the cytoplasmic domain of the type I TGF-B receptor and heterodimerization with the type II receptor are both required for intracellular signal transduction. In The EMBO Journal (pp. 4485–4496). https://doi.org/10.1007/978-3-030-24436-1_17

Massagué, J. (1998). TGFβ Signal Transduction. Annual Reviews in Biochemistry, 67, 753–791. https://doi.org/10.1097/00005768-199805001-01017

Massagué, J. (2012). TGFβ signalling in context. Nature Reviews Molecular Cell Biology, 13(10), 616–630. https://doi.org/10.1038/nrm3434

Mitchell, A., Briquez, P., Hubbell, J., & Cochran, J. (2016). Engineering growth factors for regenerative medicine applications. Acta Biomaterialia, 30, 1–12. https://doi.org/10.1016/j.actbio.2015.11.007.Engineering

Oldershaw, R. A., Baxter, M. A., Lowe, E. T., Bates, N., Grady, L. M., Soncin, F., Brison, D. R., Hardingham, T. E., & Kimber, S. J. (2010). Directed differentiation of human embryonic stem cells toward chondrocytes. Nature Biotechnology, 28(11), 1187–1194. https://doi.org/10.1038/nbt.1683

Pfaff, N., Lachmann, N., Ackermann, M., Kohlscheen, S., Brendel, C., Maetzig, T., Niemann, H., Antoniou, M. N., Grez, M., Schambach, A., Cantz, T., & Moritz, T. (2013). A ubiquitous chromatin opening element prevents transgene silencing in pluripotent stem cells and their differentiated progeny. Stem Cells, 31(3), 488–499. https://doi.org/10.1002/stem.1316

Pudasaini, A., El-Arab, K. K., & Zoltowski, B. D. (2015). LOV-based optogenetic devices: Light-driven modules to impart photoregulated control of cellular signaling. Frontiers in Molecular Biosciences, 2(MAY), 1–15. https://doi.org/10.3389/fmolb.2015.00018

Repina, N. A., Bao, X., Zimmermann, J. A., Joy, D. A., Kane, R. S., & Schaffer, D. V. (2019). Optogenetic control of Wnt signaling for modeling early embryogenic patterning with human pluripotent stem cells. BioRxiv, 665695. https://doi.org/10.1101/665695

Sako, K., Pradhan, S. J., Barone, V., Inglés-Prieto, Á., Müller, P., Ruprecht, V., Čapek, D., Galande, S., Janovjak, H., & Heisenberg, C. P. (2016). Optogenetic Control of Nodal Signaling Reveals a Temporal Pattern of Nodal Signaling Regulating Cell Fate Specification during Gastrulation. Cell Reports, 16(3), 866–877. https://doi.org/10.1016/j.celrep.2016.06.036

Salazar, V. S., Gamer, L. W., & Rosen, V. (2016). BMP signalling in skeletal development, disease and repair. Nature Reviews Endocrinology, 12(4), 203–221. https://doi.org/10.1038/nrendo.2016.12

Savary, K., Caglayan, D., Caja, L., Tzavlaki, K., Bin Nayeem, S., Bergström, T., Jiang, Y., Uhrbom, L., Forsberg-Nilsson, K., Westermark, B., Heldin, C. H., Ferletta, M., & Moustakas, A. (2013). Snail depletes the tumorigenic potential of glioblastoma. Oncogene, 32(47), 5409–5420. https://doi.org/10.1038/onc.2013.67

Tabata, Y., & Lutolf, M. P. (2017). Multiscale microenvironmental perturbation of pluripotent stem cell fate and self-organization. Scientific Reports, 7(March), 1–11. https://doi.org/10.1038/srep44711

Takahashi, F., Yamagata, D., Ishikawa, M., Fukamatsu, Y., Ogura, Y., Kasahara, M., Kiyosue, T., Kikuyama, M., Wada, M., & Kataoka, H. (2007). AUREOCHROME, a photoreceptor required for photomorphogenesis in stramenopiles. Proceedings of the National Academy of Sciences of the United States of America, 104(49), 19625–19630. https://doi.org/10.1073/pnas.0707692104

Tichy, A. M., Gerrard, E. J., Legrand, J. M. D., Hobbs, R. M., & Janovjak, H. (2019). Engineering Strategy and Vector Library for the Rapid Generation of Modular Light-Controlled Protein–Protein Interactions. Journal of Molecular Biology, 431(17), 3046–3055. https://doi.org/10.1016/j.jmb.2019.05.033

Wang, T., Nimkingratana, P., Smith, C. A., Hardingham, T. E., & Kimber, S. J. (2019). Enhanced chondrogenesis from human embryonic stem cells. Stem Cell Research, 39(February), 101497. https://doi.org/10.1016/j.scr.2019.101497

Wang, X., Chen, X., & Yang, Y. (2012). Spatiotemporal control of gene expression by a light-switchable transgene system. Nature Methods, 9(3), 266–269. https://doi.org/10.1038/nmeth.1892

Yang, G., Yuan, G., Li, X., Liu, P., Chen, Z., & Fan, M. (2014). BMP-2 induction of Dlx3 expression is mediated by p38/Smad5 signaling pathway in osteoblastic MC3T3-E1 cells. Journal of Cellular Physiology, 229(7), 943–954. https://doi.org/10.1002/jcp.24525

Ye, J., Bates, N., Soteriou, D., Grady, L., Edmond, C., Ross, A., Kerby, A., Lewis, P. A., Adeniyi, T., Wright, R., Poulton, K. V., Lowe, M., Kimber, S. J., & Brison, D. R. (2017). High quality clinical grade human embryonic stem cell lines derived from fresh discarded embryos. Stem Cell Research and Therapy, 8(1), 1–13. https://doi.org/10.1186/s13287-017-0561-y

Ziegler, T., & Möglich, A. (2015). Photoreceptor engineering. Frontiers in Molecular Biosciences, 2(JUN), 1–25. https://doi.org/10.3389/fmolb.2015.00030

